# Recurrent Neural Network Exploration Strategies During Reinforcement Learning Depend on Network Capacity

**DOI:** 10.1101/2025.03.13.642959

**Authors:** H. Flimm, D. Tuzsus, I. Pappas, J. Peters

**Affiliations:** Department of Psychology, Ludwig Maximilian University of Munich, Germany; Department of Psychology, Biological Psychology, University of Cologne, Germany; Department of Neurology, Keck School of Medicine, University of Southern California, USA

## Abstract

Artificial neural networks constitute simplified computational models of neural circuits that might help understand how the biological brain solves and represents complex tasks. Previous research revealed that recurrent neural networks (RNNs) with 48 hidden units show human-level performance in restless four-armed bandit tasks but differ from humans with respect to the task strategy employed. Here we systematically examined the impact of network capacity (no. of hidden units) on computational mechanisms and performance. Computational modeling was applied to investigate and compare network behavior between capacity levels as well as between RNNs and human learners. Using a task frequently employed in human cognitive neuroscience work as well as in animal systems neuroscience work, we show that high-capacity networks displayed increased directed exploration and attenuated random exploration relative to low-capacity networks. RNNs with 576 hidden units approached “human-like” exploration strategies, but the overall switch rate and the level of perseveration still deviated from human learners. In the context of the resource-rational framework, which posits a trade-off between reward and policy complexity, human learners may devote more resources to solving the task, albeit without performance benefits over RNNs. Taken together, this work reveals the importance of network capacity on exploration strategies during reinforcement learning and therefore contributes to the goal of building neural networks that behave “human-like” to possibly gain insights into computational mechanisms in human brains.

## Introduction

Reinforcement learning (RL) is a key theoretical account of how agents make decisions in the face of punishment or reward (Sutton & Barto, 2018). In this context, agents continuously interact with the environment, and receive environmental feedback (e.g., rewards), which then informs subsequent decisions. For example, in a typical (restless) bandit task, agents have to select among a number of options (“bandits”) with dynamically changing rewards. Computational modeling of behavior in such tasks suggests that humans use directed or strategic exploration strategies (which refers to the bias of switching to highly informative options; e.g., Chakroun et al., 2020; Wiehler et al., 2021; Wilson et al., 2014, 2021) combined with perseveration, a property that relates to the tendency to repeat previous actions independent of the experienced outcome (Badre et al., 2012; Chakroun et al., 2020; Lau & Glimcher, 2005; Payzan-LeNestour & Bossaerts, 2012; Schönberg et al., 2007; Tuzsus, Brands, et al., 2024; Wiehler et al., 2021).

Recurrent neural networks (RNNs) can be applied to RL problems due to their recurrent connectivity pattern. At each time step, RNN hidden units receive information regarding the network’s activation state at the previous time step via recurrent connections, thereby endowing the network with memory about what has happened before (Botvinick et al., 2020; Das et al., 2023; Goodfellow et al., 2016). Training and analysis of such models offer potential novel insights with implications for neuroscience (Botvinick et al., 2020). Previous studies have investigated the performance of RNNs in bandit tasks (Findling & Wyart, 2020; Tuzsus, Brands, et al., 2024; Wang et al., 2018). Recently, Tuzsus, Brands and colleagues (2024) used the restless four-armed bandit task (Daw et al., 2006) to examine the networks’ exploration strategies extensively and to compare their performance to human participants from the placebo condition of Chakroun et al. (2020). RNNs demonstrated human-level overall performance but showed no evidence of directed exploration, and higher levels of perseveration (Tuzsus, Brands, et al., 2024).

Theoretical models have suggested that perseveration simplifies the action selection and therefore constitutes “a natural consequence of limitations on policy complexity” (Gershman, 2020, p. 5). Gershman (2020) argued that resource-rational agents trade-off reward for policy complexity such that agents with limited available resources will exhibit perseveration. It follows that increasing network capacity may lead to an increase in directed exploration and reduced perseveration (Binz & Schulz, 2022; Gershman, 2020; Lai & Gershman, 2024; Meder et al., 2021; Wilson et al., 2014). Binz and Schulz (2022) provided evidence for shifts in reward-based learning strategies as a function of available computational resources. These effects resembled decision-making behavior in humans under different levels of processing capacity, including limited cognitive abilities (Gonzalez, 2004), working memory capacity (Fletcher et al., 2011), or capacity constraints attributable to brain lesions (Bechara et al., 1994) or developmental stages (Somerville et al., 2017).

Here we tested this hypothesis directly in the context of an RL task widely used to study human, animal and machine exploration behavior - the restless bandit task. We systematically explored the effects of increasing neural network capacity, i.e. the number of RNN hidden units. RNNs with varying capacity levels were trained and tested on restless four-armed bandit problems (Daw et al., 2006). Overall performance measures and model-based estimates of computational strategy were compared across capacity levels as well as between network and human learners (Chakroun et al., 2020; Tuzsus, Brands, et al., 2024) to test the hypothesis that RNNs with higher capacity would exhibit more “human-like” exploration strategies.

## Methods

The analyses for this work were conducted in R (R Core Team, 2023), JASP (JASP Team, 2024) and STAN (*rstan* package in R, Stan Development Team, 2023) to examine two research questions: first, does neural network capacity affect the performance and strategies applied in the restless four-armed bandit task? Second, do high-capacity networks approach human behavioral strategies?

### Neural Network Models

The present work used the same RNN architecture and training procedure as in Tuzsus, Brands, et al. (2024). Long short-term memory networks (LSTM; Hochreiter & Schmidhuber, 1997) with computation noise (Findling & Wyart, 2020) and no entropy regularization term (Mnih et al., 2016; Wang et al., 2018; Williams & Peng, 1991) completed 50,000 training episodes of the restless four-armed bandit task with 300 trials per episode. The network weights and biases were adjusted during training according to the advantage-actor-critic (A2C) learning algorithm (Mnih et al., 2016; Wang et al., 2017) and were held fixed during subsequent tests (see Tuzsus, Brands, et al., 2024), for details).

### Task Environment and Performance Measures

Following training, all networks were tested on the three random walk instances introduced by Daw et al. (2006), for which human data was available (*n* = 31; Chakroun et al., 2020). Task details can be found elsewhere (Daw et al., 2006; Tuzsus, Brands, et al., 2024). Performance was evaluated in terms of cumulative regret (CR). Cumulative regret is the sum of the differences between the maximum reward possible and the reward obtained in each trial of an episode, such that lower CR values reflect better performance (Agrawal & Goyal, 2012; Auer et al., 2002; Tuzsus, Brands, et al., 2024; Wang et al., 2018). In addition, the percentage of switch trials was examined, i.e. the percentage of trials in which an option different from the previous trial was selected.

### Neural network instances

Six different groups of RNNs with 30 instances each were used that differed with respect to network capacity (no. of units in the hidden layer). Capacity refers to the “ability to fit a wide variety of functions” (Goodfellow et al., 2016, pp. 111–112). We included networks with 48, 64, 80, 96, 192, or 576 hidden units, respectively. Note that the exact number of hidden units per group was selected heuristically, building on previous publications (Tuzsus, Brands, et al., 2024; Wang et al., 2018). Initially, four groups of networks (48 to 96 hidden units) were trained and tested, and two more levels were added in a subsequent step. Because two network instances in the group of networks with 576 hidden units performed extremely poorly following training (CR > 8000), they were re-trained. Furthermore, data from 31 human participants who performed the same set of task instances as the networks (one run per participant in the placebo condition) was used for comparison with human behavior (Chakroun et al., 2020; Tuzsus, Brands, et al., 2024). In total, analyses included 180 RNNs (6 groups x 30 RNN instances; 540 runs) and 31 human agents (31 runs).

### Computational Modeling

Fintz et al. (2022) proposed to use cognitive models to explain the behavior of neural networks and thereby addressing the explainability problem (e.g., Kay, 2018). Here, computational modeling was applied to gain insights into the task strategy underlying RNN performance. Two different variants of reinforcement learning models were fitted to the data. The same learning rule was used for both models (Bayesian learner). Specifically, both models were standard Kalman filter models which assume that agents track the uncertainty associated with each option, in addition to its expected outcome or value (Daw et al., 2006; Kalman, 1960; Tuzsus, Brands, et al., 2024; Wiehler et al., 2021). As opposed to the simple Rescorla-Wagner model (Rescorla & Wagner, 1972; Zhang et al., 2020), Kalman filter models incorporate an uncertainty-dependent learning rate, known as the *Kalman gain* (Daw et al., 2006; Kalman, 1960; Tuzsus, Brands, et al., 2024; Wiehler et al., 2021): learning is enhanced – represented by a higher learning rate – if uncertainty concerning the consequence of the chosen option is high compared to the situation in which the reward of the chosen option is less uncertain. Uncertainty regarding an option’s outcome decreases after sampling this option (Wiehler et al., 2021). The expected mean payoff 𝜇^ of the sampled option 𝑐_!_ is updated as follows: with the Kalman gain 𝜅_!_:

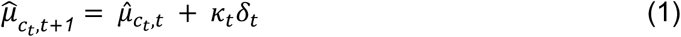

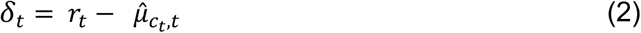

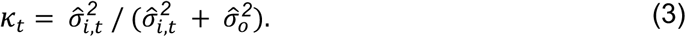

In addition, the variance of the expected mean payoff (uncertainty) is updated for the selected bandit:

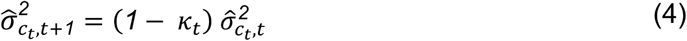

Moreover, the expected mean payoffs and uncertainties of all i bandits are updated after each trial as defined by the following equations:

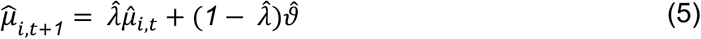

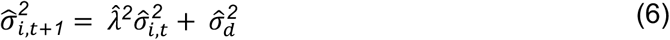

where 𝜆 denotes the decay and 𝜗 denotes the decay center. Both parameters as well as the diffusion variance 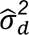 were fixed to the values from Daw and colleagues’ (2006) study 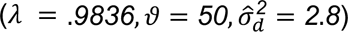.

The choice rules of both models were based on the standard softmax function (SM) but differed with regard to the number of included parameters (choice rule 5 and 12 from Tuzsus, Brands, et al., 2024). In the first model (SM + DP), three different free parameters affect the probability 𝑃 that bandit i is selected on trial t:

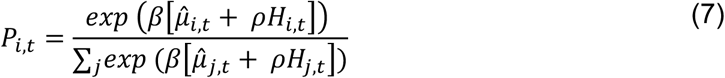

The tendency to perseverate is represented by the perseveration bonus parameter 𝜌 that assigns a bonus to the selected action from the previous trial (e.g., Chakroun et al., 2020; Tuzsus, Brands, et al., 2024; Wiehler et al., 2021). Here, it scales the degree to which the habit strength 𝐻 (Miller et al., 2019) influences the probability of selecting bandit i on trial t. The habit strength vector is computed as follows:

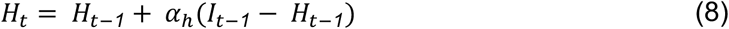

𝐻*_t_* represents a recency-weighted average of the frequency of previous actions that is updated similarly to the expected mean payoff of the selected bandit (Miller et al., 2019). The indicator function 𝐼 equals 1 for the previously selected option, and 0 otherwise (Tuzsus, Brands, et al., 2024). The free parameter 𝛼*_h_* (Miller et al., 2019; Tuzsus, Brands, et al., 2024) scales the extent to which previous choices affect the update and in turn, the choice probabilities in the current trial. It can take on values between 0 and 1. If only the previous choice influences the following choice, then 𝛼*_h_* = 1 (first-order perseveration). The lower 𝛼_*_, the higher the number of previous choices that influence the current choice (higher-order perseveration). The inverse temperature parameter 𝛽 quantifies the extent of choice stochasticity and regulates the impact of the expected mean payoff and the perseveration terms on choice probability: choices become less random (more exploitative) with increasing 𝛽 values (e.g., (Daw et al., 2006; Wiehler et al., 2021). For model fitting, 𝛽 was restricted to values ≥ 0.

The second model (SM + EDP) additionally includes the exploration bonus parameter 𝛷, a term for directed exploration (e.g., (Chakroun et al., 2020; Daw et al., 2006; Tuzsus, Brands, et al., 2024; Wiehler et al., 2021):

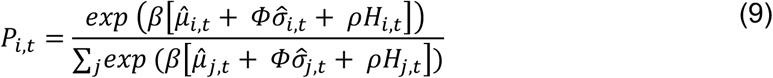

𝛷 quantifies the impact of uncertainty regarding the outcome of each choice 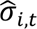. More positive values of 𝛷 reflect directed (uncertainty-based) exploration, whereas values below zero reflect uncertainty aversion (Tuzsus, Brands, et al., 2024).

Models were fitted separately for each agent with two chains and 3000 iterations per chain (1000 warm-up iterations). In total, 300 models were estimated (150 agents x 2 models = 300). Some model fits (SM + EDP) for agents with 48 hidden units and human learners were already available and used from Tuzsus, Brands, et al. (2024). The provided models were fitted to individual runs, resulting in three model fits for each RNN (30 RNN instances x 3 runs = 90 models). Fits for human participants included only a single run (= 31 models) since each participant only performed one task version in the placebo condition of Chakroun et al. (2020). The potential scale reduction factor 𝑅^ was used to evaluate chain convergence (Gelman & Rubin, 1992; Stan Development Team, 2024). Single runs were removed (*n* = 9, 2%) in case of 𝑅^ > *1*.*02* for at least one free parameter value (𝛼_*_, 𝛽, 𝜌, 𝛷).

### Model Comparison

Models were compared based on the Widely Applicable Information Criterion (WAIC), with lower estimates corresponding to a better model fit (Vehtari et al., 2017; Watanabe, 2010). WAIC values were computed separately for each run per agent. We then examined the proportion of runs for which the SM + EDP model provided a superior fit, by computing the difference in WAIC values between SM + DP and SM + EDP models per run and agent:

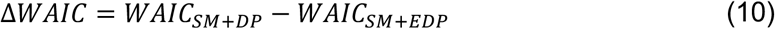

Here, Δ𝑊𝐴𝐼𝐶 > *0* indicates a superior fit for the more complex model including an exploration bonus (SM + EDP).

Model comparison only included five groups of RNNs since Tuzsus, Brands, et al. (2024) already performed a comprehensive model comparison for the group of networks with 48 hidden units as well as for the human participants from Chakroun et al. (2020). The respective fits of the selected model were provided for the comparison of model parameters in this work.

### Comparison of Model Parameters

For every agent, the median of the posterior distribution was computed for each parameter for subsequent analyses. Model parameters of the RNNs were averaged across runs to obtain a single parameter value per RNN instance. Examination of model parameters focused on the SM + EDP model, as it provided a superior fit to the behavior of human learners (Tuzsus, Brands, et al., 2024) as well as high-capacity networks – our primary groups of interest. Additionally, the SM + DP model is nested within the SM + EDP model, with both models being equal in case of 𝛷 = *0*. Therefore, the SM + EDP model contains all the information the SM + DP model contains and allowed us to directly test whether there is evidence for directed exploration (i.e. 𝛷 > *0*) (Kruschke, 2015). Due to the extensive analysis of RNNs with 48 hidden units reported in Tuzsus, Brands et al. (2024), the present model comparison only included five groups of RNNs (64, 80, 96, 192, and 576 hidden units).

### Evaluation of Capacity Effects and Differences Between RNN and Human Behavior

Differences between capacity levels in terms of model-free (cumulative regret, switch rate) and model-based measures (model parameters) were assessed using Bayesian ANOVAs (as implemented in JASP; Version 0.18.3, JASP Team, 2024; see Rouder et al., 2012, 2016, 2017; Wagenmakers et al., 2018; for details on the Bayesian ANOVA approach) for each dependent variable and with the number of hidden units as between-subjects factor. Here, evidence in favor of the alternative hypothesis indicates the existence of group differences or capacity effects, respectively. Post hoc comparisons were performed to evaluate differences between single groups. Here, the adjusted posterior odds are reported to account for multiple testing (Van Den Bergh et al., 2020; Westfall et al., 1997).

Differences between network and human behavior were evaluated using Bayesian Mann-Whitney U tests (Van Doorn et al., 2020). This comparison focused on human learners and high-capacity RNNs, which were the only network group exhibiting evidence for directed exploration (see Results).

### Associations Between Performance Measures and Model Parameters

To further understand differences in network performance, associations between performance measures (cumulative regret, switch rate) and model parameters across capacity levels were examined. Pearson correlations were calculated based on the average performance metrics and model parameters of the RNNs across runs (*N* = 180 observations per variable). In addition, associations between performance measures and model parameters were contrasted between RNNs and human learners to reveal differences in mechanisms.

### Availability of Data and Code

RNN behavioral data, R code (including Stan model code) and JASP files created for the analyses reported in this paper can be found here (view-only): https://osf.io/h76qr/?view_only=2eab7d5528e84b8da02de7d5304d7e05. Public sharing of data and code is planned after publishing the results. Human data and Python code for the training and testing of RNNs are stored in another repository (from Tuzsus, Brands, et al., 2024): https://github.com/deniztu/p1_generalization.

## Results

Networks were trained and tested on the four-armed restless bandit task previously applied in human work (Chakroun et al., 2020; Daw et al., 2006; see Methods). We investigated whether network capacity affects overall performance and underlying mechanisms in the task using computational modeling.

### Model-Agnostic Results

First, performance in terms of cumulative regret (CR) improved from the lowest (𝑀 = *1393*.*8*) to the highest network capacity (𝑀 = *1209*.*35*, Figure 1A, Table S1, note that lower values of CR reflect better performance). However, the Bayesian ANOVA did not provide evidence in favor of group differences between capacity levels (𝐵𝐹*_10_*= .*39*), and the posterior evidence for the performance improvement from 48 to 576 hidden units (𝑝𝑜𝑠𝑡𝑒𝑟𝑖𝑜𝑟 𝑜𝑑𝑑𝑠 = *1*.*61*) was inconclusive (Kurt, 2019). The group-level mean CR was lower in RNNs with 576 hidden units compared to human learners (𝑀 = *1389*.*63*), but there was again no conclusive evidence for a meaningful difference (Bayesian Mann-Whitney U Test: 𝐵𝐹*_10_* = .*4*, Figure 1C).

**Figure 1.**
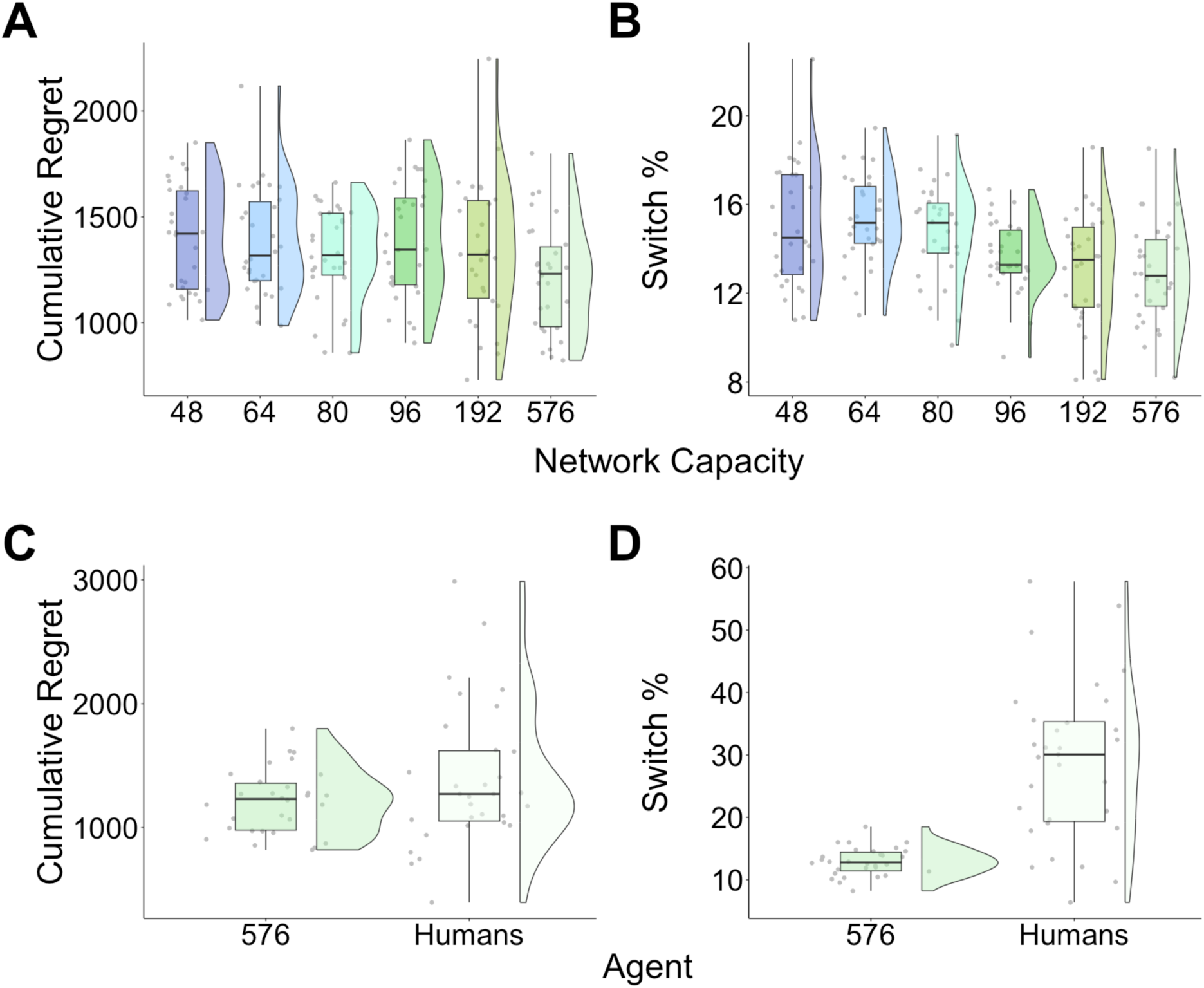
Model-agnostic results. Top panels: X-axes display network capacity in terms of the no. of hidden units in the network. Bottom panels: X-axes display agent: 576-unit networks and human learners. (A), (C) Overall performance. (B), (D) Switch rate. Each data point represents the average performance outcome for a network instance across runs.

There was an effect of capacity on switch rate (𝐵𝐹*_10_*= *1693*.*47*; Figure 1B, Table S1) and networks with 48 hidden units (𝑀 = *15%*) switched more frequently than networks with 576 hidden units (𝑀 = *12*.*9%*, 𝑝𝑜𝑠𝑡𝑒𝑟𝑖𝑜𝑟 𝑜𝑑𝑑𝑠 = *4*.*9*). Human switch rates showed a wide range (𝑀𝑖𝑛 = *6*.*4%*, 𝑀𝑎𝑥 = *57*.*8%*) and the mean switch rate of the human participants (𝑀 = *29%*) was almost twice as high as the highest average switch rate of all groups of RNNs (64 hidden units: 𝑀 = *15*.*3%*). There was strong evidence for higher switch rates in human learners compared to high-capacity RNNs (Bayesian Mann-Whitney U Test: 𝐵𝐹*_10_*= *150*.*26*; Figure 1D). Taken together, despite similar overall performances between networks and human learners, the large difference in switching behavior suggests that they may have employed, at least in part, different strategies.

### Model Comparison

Differences in model fit between the SM + DP and the SM + EDP model with an additional exploration bonus parameter were compared between groups of RNNs across capacity levels, to test the degree to which network capacity impacted directed exploration mechanisms.

The percentage of runs in which the SM + EDP model accounted best for the data increased with increasing capacity of the networks (see Figure 2). The SM + DP model provided a superior fit in more than half of the runs performed by RNNs with 64 to 192 hidden units (Figure 2 and Table S2). In contrast, the median Δ𝑊𝐴𝐼𝐶 was substantially above 0 for the 576-unit networks, indicating that incorporation of a directed exploration mechanism substantially improved model fit in these networks. For reasons listed above (see Methods), we chose the SM + EDP model for further analysis.

**Figure 2.**
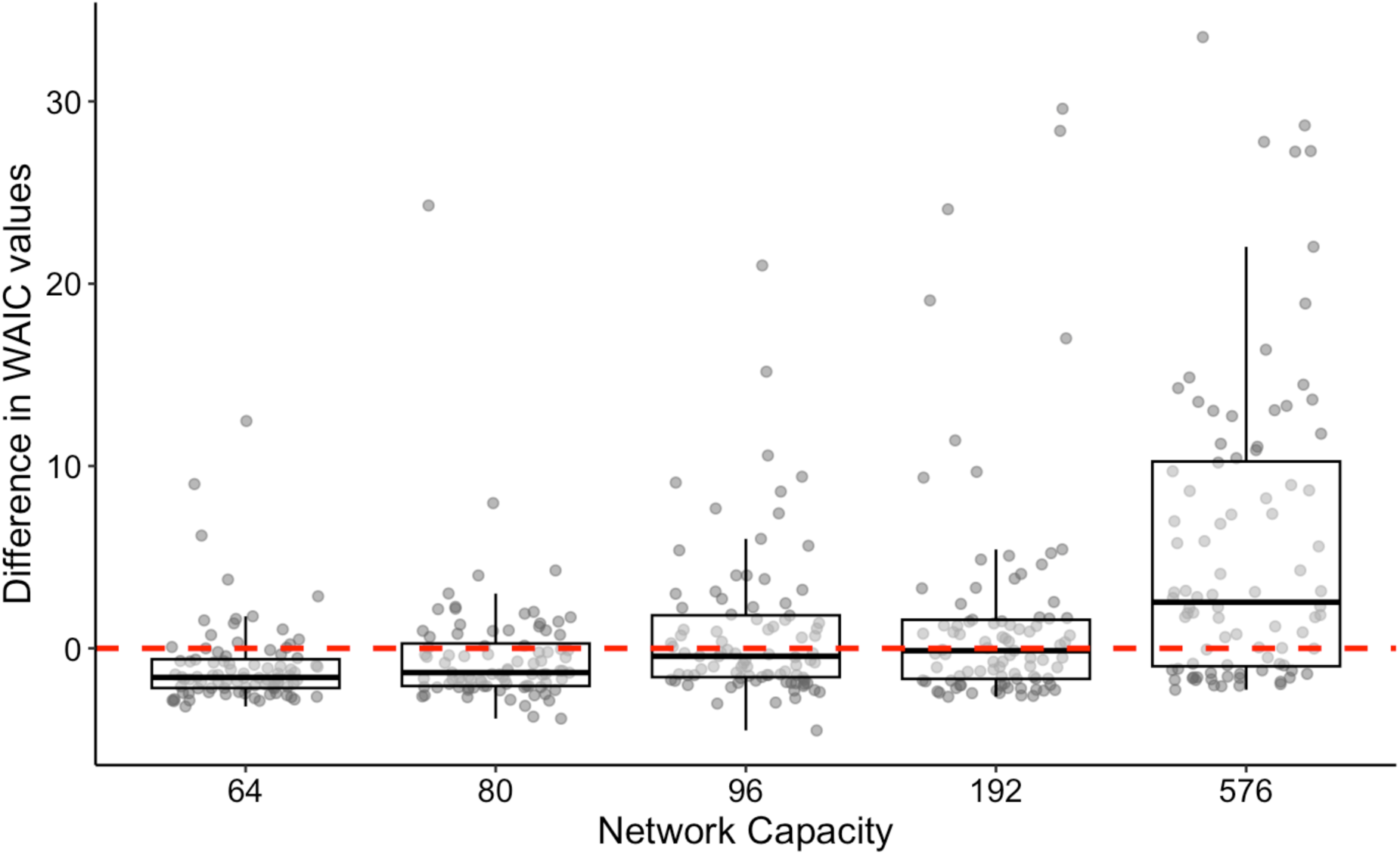
Relative Model Fit as a Function of Network Capacity. X-axes display network capacity in terms of the no. of hidden units in the network. The red dashed line indicates an equal fit of the SM + DP and SM + EDP model. Data points above the red line represent runs in which the SM + EDP model provided a superior fit to the data compared to the SM + DP model. Data points above the red line represent runs that were best accounted for by the SM + EDP model.

Figure 2 also shows that incorporating the exploration bonus parameter into the model often improved model fit substantially (Δ𝑊𝐴𝐼𝐶 > 0) for networks with a smaller number of units. In contrast, when the SM + DP model accounted better for the data, the differences in model fit were numerically small in all cases (Δ𝑊𝐴𝐼𝐶 < *0*, but close to 0). In simpler terms, when the SM + DP model accounted best for the data (data points below the red line), the SM + EDP model provided only a marginally inferior fit (data points near 0). Conversely, when the SM + EDP model accounted best for the data (data points above the red line), the SM + DP model frequently yielded a substantially poorer fit (data points farther from 0).

### Comparison of Model Parameters

We next compared model parameters across networks with different capacity levels and between high-capacity networks and human learners to quantify similarities and differences in exploration and perseveration behavior.

Consistent with the model comparison results, directed exploration was modulated by network capacity: increasing capacity resulted in increased directed exploration (Table S3, 𝐵𝐹*_10_*= *5*.*84* × *10^10^*). RNNs with 576 hidden units displayed the highest level of directed exploration (𝑀 = .*802*). Human learners exhibited numerically higher levels of directed exploration (𝑀 = *1*.*479*) compared to 576-unit networks, but the Bayesian Mann-Whitney U test was inconclusive (𝐵𝐹*_10_*= .*97*; Figure 3C).

**Figure 3.**
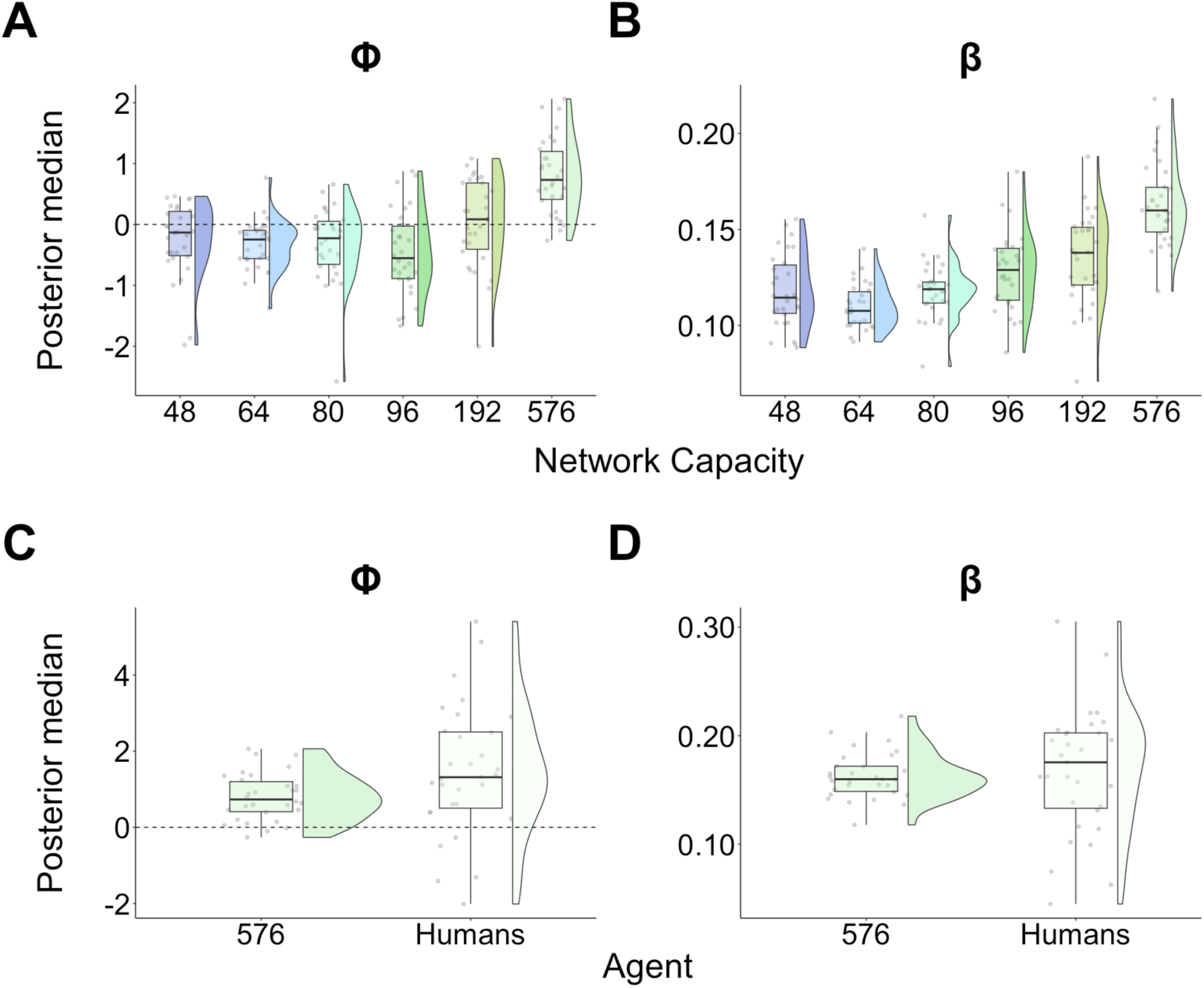
Exploration behavior. Top panels: X-axes display network capacity in terms of the no. of hidden units in the network. Bottom panels: X-axes display agent: 576-unit networks and human learners. (A), (C) Φ: Exploration bonus. (B), (D) β: Inverse temperature parameter. RNN data was aggregated across runs to obtain a single value for each network instance (mean).

In addition, we found that the higher the capacity of the networks, the lower the degree of random exploration as indicated by the increase in the inverse temperature parameter (𝐵𝐹*_10_*= *2*.*62* × *10^19^*, Figure 3B; Table S3, two extreme outliers in the group of RNNs with 80 hidden units had been removed for the Bayesian ANOVA). While posterior evidence suggested that the inverse temperature parameter 𝛽 increased from RNNs with 48 hidden units (𝑀 = .*118*) to RNNs with 576 hidden units (𝑀 = .*163*; 𝑝𝑜𝑠𝑡𝑒𝑟𝑖𝑜𝑟 𝑜𝑑𝑑𝑠 = *2*.*8* × *10^8^*), the degree of random exploration was comparable between high-capacity networks and human learners (𝑀 = .*168*; Bayesian Mann-Whitney U test: 𝐵𝐹*_10_*= .*31*; Figure 3D).

Perseveration levels remained relatively constant across network capacities (𝐵𝐹*_10_*= *2*.*65*, Figure S1A). The level of perseveration displayed by 576-unit networks (𝑀 = *15*.*86*) exceeded human-level perseveration (𝑀 = *10*.*9*, Bayesian Mann-Whitney U test: 𝐵𝐹*_10_*= *15*.*48*; Figure S1C). Results further demonstrate that the extent of higher-order perseveration decreases as the capacity of the networks increases (𝐵𝐹*_10_* = *2*.*32* × *10^8^*, Figure S1B). Here, the mean parameter value of 𝛼_*_ for human learners (𝑀 = .*57*) was similar to the group of RNNs with the lowest capacity (𝑀 = .*65*). The Bayesian Mann-Whitney U test yielded convincing evidence in favor of differences in higher-order perseveration between RNNs with 576 hidden units and human learners (𝐵𝐹*_10_*= *109*.*36*, Figure S1D).

Consequently, increasing the number of hidden units of the networks did not lead to human-like perseveration behavior. However, we observed a reduction in perseveration with an increasing number of hidden units that is characterized by a reduction in higher-order perseveration, rather than a decrease in general perseveration tendencies.

### Associations Between Performance Measures and Model Parameters

Having identified differences in exploration and perseveration as a function of capacity (see above), we sought to understand how these relate to overall task performance. We performed correlations between performance measures and model parameters, and we found that a higher switch rate was associated with worse performance for both humans (𝑟(*29*) = .*56*, 𝑝 = .*001*) and RNNs (𝑟(*178*) = .*32*, 𝑝 < .*001*). In addition, switch rate was positively associated with directed exploration in humans (𝑟(*29*) = .*45*, 𝑝 = .*012*). Consequently, humans who displayed a high level of directed exploration switched more and in turn performed worse than human learners who only moderately explored uncertain options (Figure 4B). This “over-exploration” of uncertain options appears to result in performance impairments comparable to those associated with uncertainty aversion (𝛷 < *0*). For RNNs, the switch rate (𝑟(*178*) = −.*16*, 𝑝 = .*031*) as well as cumulative regret decreased with an increasing exploration bonus parameter (𝑟(*178*) = −.*3*, 𝑝 < .*001*; Figure 4A). Therefore, increasing network capacity seems to result in more efficient switching (less *and* more directed).

**Figure 4.**
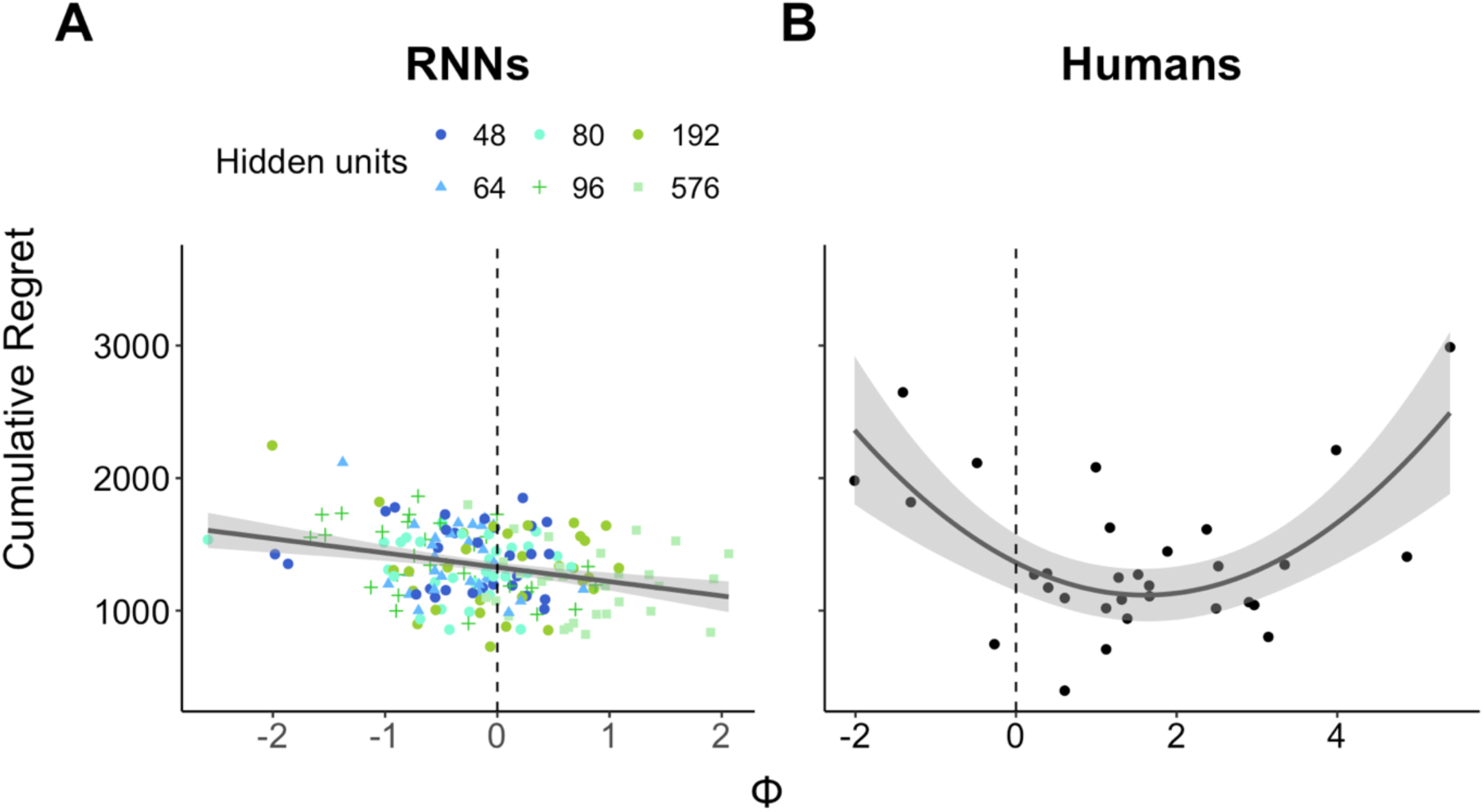
Association Between Overall Performance and Directed Exploration in RNNs vs. Human Learners. 𝛷 = Exploration bonus parameter. (A) Grey line denotes the relationship between directed exploration and overall performance across all groups of RNNs (color-coded). Data points represent the average regret of each RNN across runs. (B) Grey line denotes the relationship between directed exploration and overall performance in the group of human learners. Data points represent the regret of each participant in a single run (placebo condition; Chakroun et al., 2020).

In terms of random exploration, agents who displayed lower choice stochasticity performed better overall (RNNs: 𝑟(*176*) = −.*36*, 𝑝 < .*001*; humans: 𝑟(*29*) = −.*68*, 𝑝 < .*001*) and switched less frequently (RNNs: 𝑟(*176*) = −.*5*, 𝑝 < .*001*; humans: 𝑟(*29*) = −.*51*, 𝑝 = .*004*). On the contrary, the perseveration bonus parameter 𝜌 did not correlate with overall performance (RNNs: 𝑟(*178*) = .*14*, 𝑝 = .*068*; humans: 𝑟(*29*) = .*22*, 𝑝 = .*23*), or the percentage of switch trials (RNNs: 𝑟(*178*) ≈ *0*, 𝑝 ≈ *1*; humans: 𝑟(*29*) = .*17*, 𝑝 = .*36*). With respect to the RNNs, there was no association between 𝛼_*_ and cumulative regret (𝑟(*178*) = −.*07*, 𝑝 = .*33*) while the switch rate was inversely correlated with the extent of higher-order perseveration (𝑟(*178*) = −.*25*, 𝑝 < .*001*).

## Discussion

Here we show that in restless multi-armed bandit tasks previously applied in human work to study exploration behavior (Chakroun et al., 2020; Daw et al., 2006), the task strategy employed by RNNs depends on their respective network capacity. Networks with high capacity levels exhibited higher levels of directed exploration and attenuated random exploration, which was also reflected in a lower switch rate. This contrasts with low capacity networks, which showed no evidence of directed exploration, extending the results from Tuzsus, Brands et al. (2024). These changes in exploration behavior were associated with improved performance. Furthermore, RNNs with high capacity showed a lower degree of higher-order perseveration compared to low-capacity RNNs.

The finding of increased directed exploration in high-capacity RNNs is in line with the work of Binz and Schulz (2022) who reported an increase in directed exploration in models with greater description lengths, relative to models with shorter description lengths. Moreover, they discovered that shorter description lengths led to behavior that resembled that of patients with brain lesions from Bechara and colleagues’ study (1994). Somerville et al. (2017) further found that levels of directed exploration in humans increased during adolescence. Models with shorter or longer description lengths displayed differences in exploration strategies that corresponded to these age-related effects in humans (Binz & Schulz, 2022). Together with the results of the present work, findings suggest that the use of directed exploration depends on computational resources.

High-capacity networks showed comparable overall performance to human learners (Chakroun et al., 2020; Tuzsus, Brands, et al., 2024) and employed human-like exploration strategies. However, perseveration behavior did not approach human-level perseveration and human learners displayed significantly higher switch rates than networks. In light of the notion of resource-rational agents that trade-off reward for policy complexity (Gershman, 2020; Lai & Gershman, 2024), human behavior does not appear resource-rational since human learners did not outperform networks, although applying more demanding strategies (i.e. lower levels of perseveration and higher switch rates). If reward maximization was their only objective, humans seem to invest more resources and take more risk than necessary.

The task environment may play a role. In the task environments used here, actions were independent from each other (Daw et al., 2006), that is the reward obtained from sampling a bandit does not inform the agent about the rewards of other options. Consequently, the agents cannot form predictions that could guide their choices as in structured environments, in which the options are associated with each other (Gershman & Niv, 2010, 2015; Schulz et al., 2020). For example, Gershman and Niv (2015) instructed their participants to choose between a reference option with a known reward probability and a novel option in each trial, with the novel option either drawn from a context with a high or a low reward probability. The probability of choosing the novel option was context-dependent, implying that the participants acquired the task structure and inferred the value of unknown options on the basis of previously experienced rewards from the same context (Gershman & Niv, 2015). The only way to learn about the reward of a bandit in the unstructured task used in the present work is to select it or in other words, explore it (Daw et al., 2006). It has been stated that real-world environments typically contain structure and that the task of choosing between independent actions therefore is unrealistic (Gershman & Niv, 2010; Schulz et al., 2020). Humans might have switched so often to find some kind of pattern to base their future choices on, because it narrows the scope of what needs to be learned (Gershman & Niv, 2010; Lai & Gershman, 2024). From this perspective, human switching does have a rational component.

Furthermore, curiosity might be another reason for higher switch rates in humans. Dubey and Griffiths (2020) propose that humans are curious about stimuli that increase their current knowledge. According to their theory, exploring these stimuli is rational as knowledge about the stimuli is the key to reward maximization in the future. In an environment in which the occurrence of a stimulus does not depend on the past^1^, it would be rational to explore unknown stimuli. Curiosity reflects the underlying mechanism that motivates the agent to do so (Dubey & Griffiths, 2020). However, in the case of the bandit problem, both exploration and exploitation are necessary to maximize rewards over time (Dubey & Griffiths, 2020; Sutton & Barto, 2018). There are studies indicating that human curiosity may be able to “disturb” the balance between exploration and exploitation in reward-based learning tasks, or more generally, as Brändle et al. (2020) describe it: “there are cases where curiosity seemingly goes awry” (p. 2). For example, Gershman and Niv (2015) discovered that their participants expressed a novelty bias: they over-explored the novel option from the context with a low reward probability. Albeit previous experience suggested the selection of the reference option with a reward probability of 50% instead of the novel option with a reward probability of only 20%, participants chose the novel option in more than 40% of the trials (Gershman & Niv, 2015). The causes of suboptimal manifestations of human curiosity require further investigation (Brändle et al., 2020). Implementing curiosity in artificial agents via curiosity-driven algorithms, as has already been proposed (Ruggeri et al., 2024; Schmidhuber, 2010), might lead to a higher exploration rate and a lower level of perseveration.

A limitation of the present study regards the size of the human sample and the fact that data for each human agent comprised only one run (placebo condition of Chakroun et al., 2020). Lai and Gershman (2024) reported that humans seek to apply simple policies at the cost of performance. Here, human learners exhibited a more complex policy than networks but with no performance advantage. In future studies, it would be interesting to investigate whether human learners are able to adapt their policy to the task environment with practice. Perhaps, agents learn to reduce the complexity of their policy after acquiring more knowledge about the nature of the task. Recent work showed that human participants were able to improve their performance in a reward-based learning task with practice and that learning parameters stabilized after multiple practice runs, questioning whether parameters assessed in the first few runs are actually reliable (Neuser et al., 2023).

Furthermore, it is important to note that the task instances on which the networks were trained differed to the testing environments with respect to the diffusion variance (training: 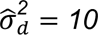, test: 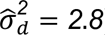 . That is, networks were trained in environments of higher volatility. Since recent findings suggest that humans reduce the complexity of their policy under higher cognitive demands (Lai & Gershman, 2024), it could be expected that humans would reduce policy complexity under higher volatility since it is harder to keep track of the current rewards of the options if they change faster or more drastically, respectively. The simplification of the policy in turn may result in a greater alignment between network and human behavior in more complex environments, e.g., humans might perseverate more (Lai & Gershman, 2024). However, we argue that higher levels of perseveration observed in RNNs relative to human learners cannot be attributed solely to the higher volatility of the training environments compared to the testing environments. First, a previous study showed that training the networks on environments with medium or varying volatility levels improved adaptation to different volatilities in the testing phase relative to a training regime where networks only encountered low-volatility environments (Tuzsus, Pappas, et al., 2024). Therefore, it is reasonable to assume that networks were able to adapt to the lower volatility of the testing environments here. Second, human learners have probably encountered environments of different volatilities throughout their lives. Thus, it is reasonable to posit that they are also capable of adapting to specific environments. Future studies might examine human and RNN policy adaptivity by contrasting behavior in environments of varying volatilities.

## Conclusions

Here we show that exploration strategies of recurrent neural networks during a restless multi-armed bandit reinforcement learning task depend on network capacity. High-capacity networks applied more sophisticated strategies as expressed by an increase in directed exploration and a reduction in random exploration relative to low-capacity networks. These shifts are consistent with the notion of more “human-like” behavior. However, high-capacity networks still deviated from human learners in terms of first-and higher-order perseveration. The comparison to human learners also suggests that networks behave more resource-rational than humans who – despite comparable overall performance – perseverate less and switch more. To conclude, this work reveals the importance of network capacity on computational mechanisms during reinforcement learning and therefore contributes to the goal of building neural networks that behave human-like (Momennejad, 2022) to possibly gain insights into computational mechanisms in human brains (Kriegeskorte, 2015; Kriegeskorte & Mok, 2017; Song et al., 2017).

## Acknowledgements

This work was funded by Deutsche Forschungsgemeinschaft (PE1627/8-1, Code 496990750 to J.P.).

## Supplemental Material

**Table S1.** Group-Level Performance Measures.

| Agents | Cumulative Regret |  | Switch % |  |
| --- | --- | --- | --- | --- |
|  | *M* | *SE* | *M* | *SE* |
| 48 | 1393.802 | 46.143 | 15.022 | 0.505 |
| 64 | 1378.122 | 46.651 | 15.346 | 0.357 |
| 80 | 1323.295 | 41.295 | 14.741 | 0.391 |
| 96 | 1382.994 | 48.856 | 13.648 | 0.291 |
| 192 | 1333.505 | 59.913 | 13.046 | 0.47 |
| 576 | 1209.349 | 47.249 | 12.895 | 0.41 |
| Humans | 1389.628 | 102.528 | 28.965 | 2.287 |

*Note.* RNNs: *N* = 30 observations per group. Humans: *N* = 31 observations.

**Table S2.** Number and Percentage of Runs in Which the SM + EDP Model Accounted Best for the Behavior of the RNNs as a Function of Network Capacity.

| No. of hidden units | $\Delta WAIC > 0$ | |
| --- | --- | --- |
|  | *n* | % |
| 64 | 16 | 18.2 |
| 80 | 25 | 28.4 |
| 96 | 37 | 42 |
| 192 | 43 | 48.3 |
| 576 | 59 | 67 |

*Note.* $\Delta WAIC$ = Difference in WAIC values between the SM + DP and the SM + EDP model; $\Delta WAIC > 0$ indicates a superior fit of the SM + EDP model over the SM + DP model.

**Table S3.** Group-Level Model Parameters.

| Agents | $\beta$ | | $\Phi$ | | $\rho$ | | $\alpha_h$ | |
| --- | --- | --- | --- | --- | --- | --- | --- | --- |
|  | *M* | *SE* | *M* | *SE* | *M* | *SE* | *M* | *SE* |
| 48 | 0.118 | 0.003 | -0.25 | 0.112 | 16.627 | 0.554 | .647 | .015 |
| 64 | 0.11 | 0.002 | -0.334 | 0.075 | 16.557 | 0.517 | .678 | .017 |
| 80 | 0.118<sup>2</sup> | 0.002 | -0.318 | 0.113 | 15.483 | 0.622 | .67 | .017 |
| 96 | 0.128 | 0.004 | -0.462 | 0.126 | 13.543 | 0.683 | .697 | .017 |
| 192 | 0.135 | 0.004 | 0.035 | 0.132 | 15.398 | 0.772 | .794 | .019 |
| 576 | 0.163 | 0.004 | 0.802 | 0.113 | 15.86 | 0.675 | .789 | .019 |
| Humans | 0.168 | 0.01 | 1.479 | 0.308 | 10.904 | 1.097 | .571 | .036 |

*Note.* *RNNs*: $N = 30$ agents per group. *Humans*: $N = 31$ agents. $\beta$ : Inverse temperature parameter; $\Phi$ : Exploration bonus; $\rho$ : Perseveration bonus; $\alpha_h$ : Habit step-size parameter.
<sup>2</sup> $n = 28$ observations

**Figure S1.**
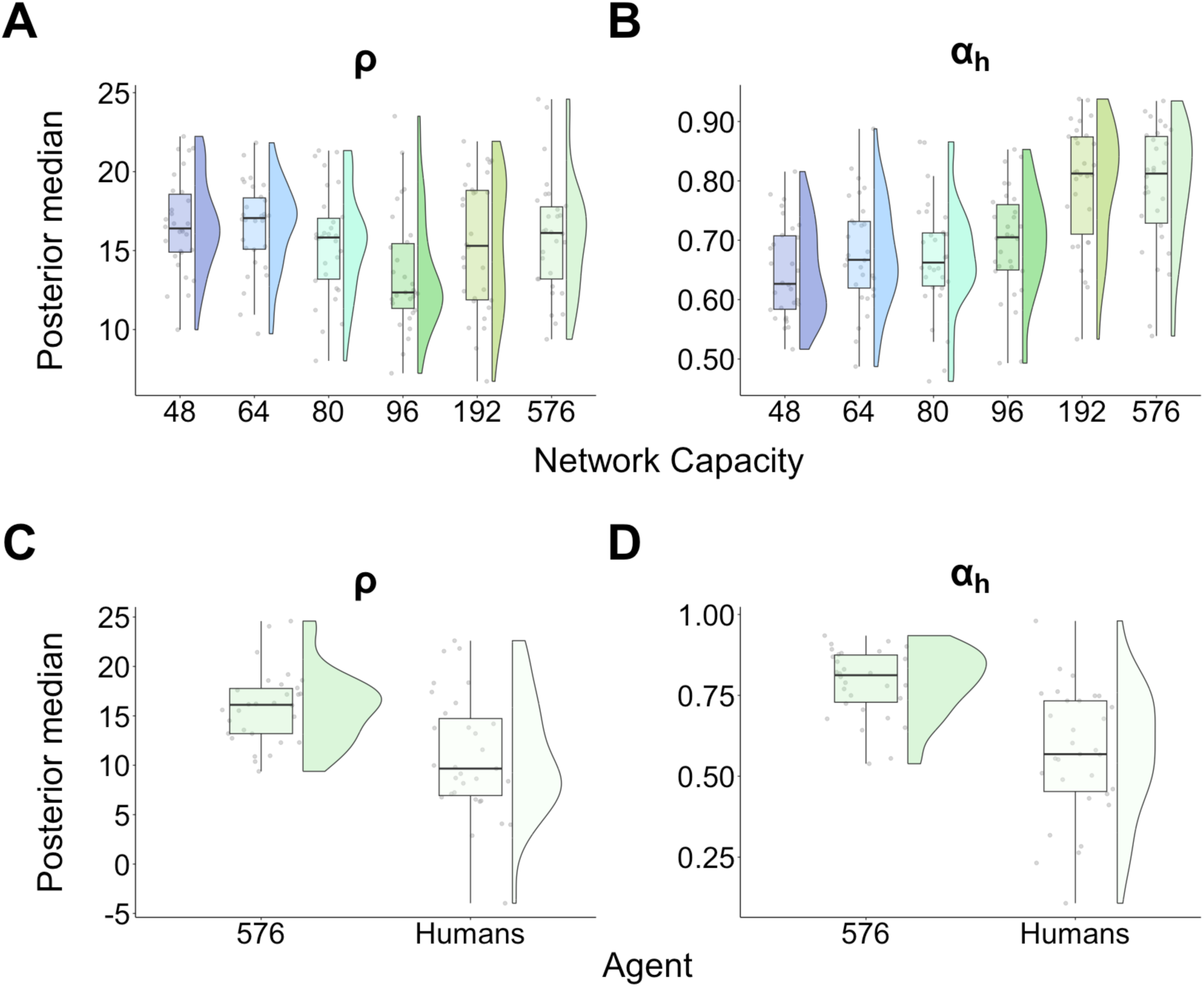
Perseveration Behavior. Top panels: X-axes display network capacity in terms of the no. of hidden units in the network. Bottom panels: X-axes display agent: 576-unit networks and human learners. (A), (C) ρ: Perseveration bonus. (B), (D) 𝛼_*_: Habit step-size parameter. RNN data was aggregated across runs to obtain a single value for each network instance (mean).

## Footnotes

1 This is the case here: in each trial, the agents can choose between the four bandits, regardless of their choices in the past (Daw et al., 2006).

